# Reduced Influence of Perceptual Context in Schizophrenia: Behavioral and Neurophysiological Evidence

**DOI:** 10.1101/795534

**Authors:** Victor J. Pokorny, Timothy J. Lano, Michael-Paul Schallmo, Cheryl A. Olman, Scott R. Sponheim

**Affiliations:** Minneapolis Veterans Affairs Health Care System; University of Minnesota Department of Psychiatry and Behavioral Science; University of Minnesota Department of Psychology

**Keywords:** Schizophrenia, Psychosis, Context, Contour, Perception, EEG

## Abstract

**Background:** Accurate perception of visual contours is essential for seeing and differentiating objects in the environment. Both the ability to detect visual contours and the influence of perceptual context created by surrounding stimuli are diminished in people with schizophrenia. The central aim of the present study was to better understand the biological underpinnings of impaired contour integration and weakened effects of perceptual context. Additionally, we sought to determine whether visual perceptual abnormalities reflect genetic factors in schizophrenia and are present in other severe mental disorders.

**Methods:** We examined behavioral data and event-related potentials (ERPs) collected during the perception of simple linear contours embedded in similar background stimuli in 27 patients with schizophrenia (SCZ), 23 patients with bipolar disorder, 23 first-degree relatives of SCZ and 37 controls.

**Results:** SCZ exhibited impaired visual contour detection while patients with bipolar disorder exhibited intermediate performance. The orientation of neighboring stimuli (i.e., flankers) relative to the contour modulated perception across all groups, but SCZ exhibited weakened suppression by the perceptual context created by flankers. Late visual (occipital P2) and cognitive (centroparietal P3) neural responses showed group differences and flanker orientation effects, unlike earlier ERPs (occipital P1 and N1). Moreover, behavioral effects of flanker context on contour perception were correlated with modulation in P2/P3 amplitudes.

**Conclusion:** In addition to replicating and extending findings of abnormal contour integration and visual context modulation in SCZ, we provide novel evidence that abnormal use of perceptual context is associated with higher-order sensory and cognitive processes.

## Introduction

Schizophrenia is a heterogeneous disorder often associated with abnormal sensory processing (O’Donnell *et al.* 2004; Butler *et al.* 2008). Aberrant processing of visual stimuli can lead to perceptual abnormalities that may manifest as visual distortions or hallucinations. Recent investigations suggest that abnormal visual processing is due in part to deviant functions of the visual cortex (Silverstein *et al.* 2015a) and may reflect abnormal sensory gain control in schizophrenia (Butler *et al.* 2008; Earls *et al.* 2016; Grove *et al.* 2018) and mark genetic liability for the disorder (Sponheim *et al.* 2013; Earls *et al.* 2016; Grove *et al.* 2018).

The detection of contours that typically define the boundaries of objects (e.g., object vs. background) is an important perceptual function for navigating the visual world. A special type of contour detection involves inferentially perceiving discrete, spatially separated edge elements as a larger, single contour, in accordance with Gestalt principles of proximity and good continuation (Wertheimer 1938). There is evidence that patients with schizophrenia (SCZ) exhibit impaired contour integration (i.e., impaired ability to perceive a spatially separated contour as a single unit, or diminished perception of shapes defined by such contours) as compared to healthy controls (Silverstein & Keane 2011); however, less is known about the neurophysiological correlates of this abnormality. To date, researchers have examined the perception of contours that form closed objects yielding evidence of abnormalities in both early visual cortical and higher-level functions in schizophrenia (Silverstein *et al.* 2000; Spencer *et al.* 2003, 2004; Foxe *et al.* 2005; Butler *et al.* 2013), but it remains unclear whether perception of simple discrete contours devoid of semantic information may yield similar high-level abnormalities. Demonstrating that simple contours produce high-level deficits in schizophrenia would provide evidence for wide-ranging effects of perceptual deficits.

Another important perceptual function for navigating the visual world involves the modulation of a central stimulus by the presence and configuration of surrounding stimuli; typically, more similar surrounding contexts reduce perceptual salience (i.e., surround suppression; (Chubb *et al.* 1989; Snowden & Hammett 1998; Xing & Heeger 2000, 2001; Yu *et al.* 2001, 2003)). Effects of surrounding context are thought to be weaker in SCZ; for example, surround suppression during contrast perception is weaker in SCZ vs. healthy control subjects (Dakin *et al.* 2005; Yoon *et al.* 2009; Tibber *et al.* 2013; Yang *et al.* 2013a; Schallmo *et al.* 2015), and similar effects have been reported during contour detection (Schallmo *et al.* 2013).

Visual contextual modulation is thought to depend on lower and higher-level processes, based on work in both animal models (Webb *et al.* 2005; Angelucci & Bressloff 2006; Nurminen *et al.* 2018) and humans (Cai *et al.* 2008; Petrov & McKee 2009; Schallmo & Murray 2016; Schallmo *et al.* 2019). Generally, lower level mechanisms (e.g., edge detection) are thought to manifest earlier in visual processing while higher level processes (e.g., shape integration) are thought to occur later in time. Few, if any, studies have delineated how psychotic psychopathology may affect these lower and higher level processes. Given the distinct time courses of these proposed mechanisms, the high temporal resolution of electroencephalography (EEG) may be useful for identifying when, during visual processing, deviant contextual modulation occurs.

To date, the extent to which impaired contour integration and weaker context modulation are specific to patients with schizophrenia, as compared to patients with bipolar disorder (BP) is unclear (Yang *et al.* 2013b; Schallmo *et al.* 2015). Schallmo and colleagues (24) reported weakened surround suppression effects in both schizophrenia and bipolar disorders as compared to controls (CON), although SCZ exhibited more profound deficits than BP. In contrast, Yang and colleagues (Yang *et al.* 2013b) found that effects of surrounding context were not significantly weaker for BP vs. CON participants, with only SCZ exhibiting weakened surround suppression. As such, more evidence is needed to clarify the relationship between weakened surround suppression and BP. It is worth noting that neither of the aforementioned studies involved contour integration and that the impact of BP on contour perception is not well understood.

There are also relatively few studies comparing contextual modulation in SCZ to first degree relatives of patients with schizophrenia (SREL; (Schallmo *et al.* 2013, 2015)). Both of these studies from our group reported normal surround suppression in SREL, but did not collect electrophysiological data and thus could not speak to the latent physiology of contextual modulation in SREL. Investigation of the neurobiology of normal contextual modulation in SREL may provide insights into the relationship between genetic liability for schizophrenia and abnormal contextual modulation.

The central aim of the present study was to identify neural correlates of impaired contextual processing and contour integration in SCZ, BP and SREL as compared to CON. More specifically, we measured ERPs in order to understand the time course of neural responses to visual contour stimuli, the modulation by surrounding context, and deficient neural processing in SCZ. Our secondary aim was to clarify the degree to which contextual modulation and contour integration are aberrant in BP and SREL to better understand the specificity of abnormal visuoperceptual processing in SCZ and genetic liability for the disorder.

## Methods and Materials

### Participants

27 SCZ, 23 SREL, 23 BP, and 37 CON completed the Collinear Gabor Contour Task (Schumacher *et al.* 2011; Schallmo *et al.* 2013) as part of two family studies of severe psychopathology at the Minneapolis VA Medical Center. Of these participants, 14 SCZ, 8 SREL and 15 CON participated in a previous study that implemented an earlier version of the Collinear Gabor Contour Task (Schallmo *et al.* 2013). Table 1 provides relevant demographic and clinical characteristics of participants. Patients were recruited from Minneapolis Veterans Affairs (VA) Medical Center outpatient clinics, community support programs for the mentally ill, and county mental health clinics. SREL were identified by research staff using a pedigree form completed through interviews with SCZ. CON were recruited via posted announcements at fitness centers, community libraries, the Minneapolis VA Medical Center, and newsletters for veterans and fraternal organizations.

**Table 1.** Participant Demographic Characteristics and Symptom Ratings

| Index | SCZ (n=27) | BP (n=23) | SREL (n=23) | CON (n=37) | Statistics |
| --- | --- | --- | --- | --- | --- |
| Age | 43.3 (10.0) | 46.1 (11.1) | 45.4 (10.6) | 47.1 (11.4) | $F(3,106) = .66, p = .58$ |
| Percent Female | 15%<sup>a</sup> | 30% | 57% | 36% | $\chi^2_{(3)} = 9.82, p = .02$ |
| Education | 13.8 (1.9) | 13.8 (1.8) | 14.9 (2.5) | 15.1 (1.8) | $F_{(3,106)} = 3.22, p = .026$ |
| Estimated IQ (from WAIS-III) | 91.3 (20.0)<sup>b</sup> | 97.9 (14.5) | 102.5 (16.1) | 103.5 (14.5) | $F_{(3,104)} = 3.27, p = .024$ |
| Overall Symptomatology (BPRS Total Score) | 44.2 (12.0)<sup>c</sup> | 36.3 (8.6)<sup>d</sup> | 34.2 (8.0) | 28.4 (4.1) | $F_{(3,106)} = 18.87, p < .001$ |
| Schizotypal Characteristics (SPQ Total Score) | 24.2 (17.3)<sup>e</sup> | 24.4 (18.5)<sup>e</sup> | 7.7 (10.9) | 6.6 (6.3) | $F_{(3,93)} = 17.67, p < .001$ |
| Abnormal Perceptual Gating (SGI Total Score) | 69 (39.2)<sup>f</sup> | 59.8 (32.4)<sup>f</sup> | 44.3 (34.8) | 34 (22.6) | $F_{(3,90)} = 6.26, p = .001$ |

All data are presented as Mean (Standard Deviation), unless otherwise noted. SCZ = patients with schizophrenia, BP = patients with bipolar disorder, SREL = first degree relatives of SCZ, CON = healthy controls. WAIS-III = Wechsler Adult Intelligence Scale, 3rd edition. BPRS = 24-item brief psychiatric Rating Scale. SPQ = Schizotypal Personality Questionnaire. SGI = Sensory Gating Inventory.
- a. There was a significant difference in gender distribution for SCZ compared to SREL, FDR corrected $p = .01$ . b. Estimated IQ was significantly lower in SCZ as compared to CON, Tukey's HSD, $p = .02$ . c. BPRS scores were higher in SCZ as compared to BP, SREL and CON, Tukey's HSD, $p s < .006$ . d. BPRS scores for BP were significantly higher than CON, Tukey's HSD, $p = .003$ . e. SPQ scores were higher for both patient groups as compared to CON and SREL, Tukey's HSD, $p s < .001$ . f. SGI scores were higher for both patient groups as compared to CON, Tukey's HSD, $p s < .03$ .
Estimated IQ data were not obtained for one patient with bipolar disorder and one healthy control. SPQ data were not obtained for one patient with schizophrenia, three SREL, four BP, and five CON. SGI data were not obtained for two SREL, three CON, five BP, and six SCZ.

Exclusion information can be found in the supplemental materials. The study protocol was reviewed, approved, and monitored by the Minneapolis VA Medical Center and the University of Minnesota Institutional Review Board. Structured Clinical Interview for the DSM-IV-TR Axis-I Disorders-Patient Edition (SCID-I/P; (First *et al.* 2002), the Brief Psychiatric Rating Scale, 24-item (BPRS; (Overall & Gorham 1962), Schizotypal Personality Questionnaire (Raine 1991), Sensory Gating Inventory (Hetrick *et al.* 2012) and the Wechsler Adult Intelligence Scale, Third Edition (WAIS-III; (Wechsler 1997) were administered to all participants. A minimum of two trained psychologists (advanced doctoral students in clinical psychology, postdoctoral researchers, or licensed psychologists) reached consensus on all diagnoses, based on the *DSM-IV-TR* criteria (American Psychological Association 2000). Additional participant and study information is detailed in previous publications (Goghari *et al.* 2014; Lynn *et al.* 2016).

### Participant Characteristics

Participant demographic information is presented in Table 1. Due to differences in gender distribution across groups gender was added as a between-subjects factor in all analyses (Steffensen *et al.* 2008; Feng *et al.* 2011). There were differences in education across groups, but follow-up pairwise comparisons between groups did not reach significance. Estimated IQ differed across groups with CON exhibiting higher IQs than SCZ. Psychiatric symptoms, as assessed by the BPRS, were more prominent in SCZ as compared to BP, CON, and SREL while BP exhibited intermediate psychiatric symptomatology, scoring lower than SCZ, but higher than CON. Schizotypal personality trait (SPQ) scores were higher for both patient groups as compared to SREL and CON. Similarly, phenomenological perceptual gating abnormalities, as assessed by the SGI, were more prominent in both patient groups as compared to SREL and CON.

### Stimuli

Stimuli details can be found in the supplemental materials. Target contours were made up of five vertically aligned Gabor patches centered at 1.6° eccentricity to the left or right of a central fixation point along the horizontal meridian (see Figure 1). Target contour detection thresholds were calculated for three flanker conditions: random, parallel, and orthogonal.

**Figure 1.**
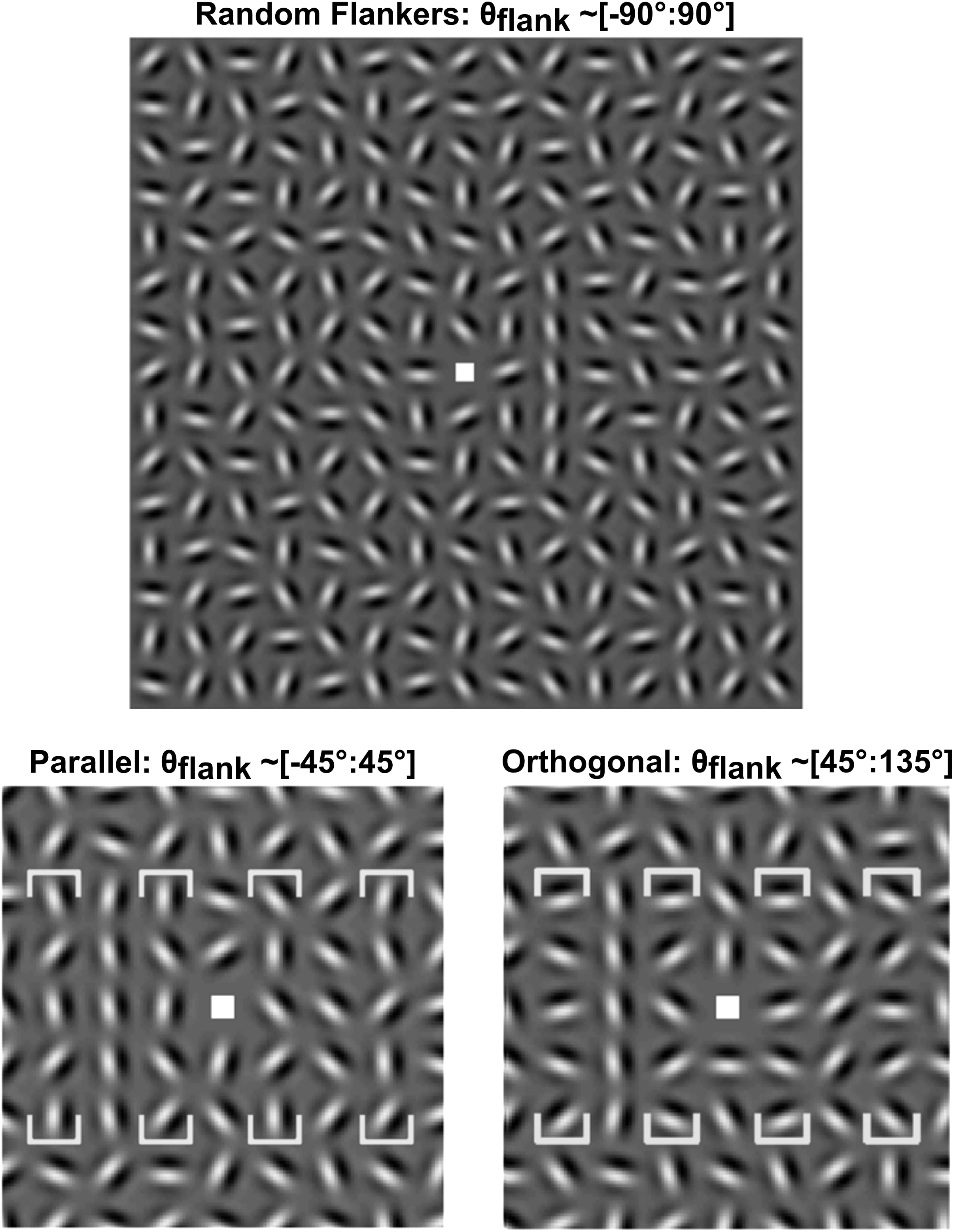
Stimulus Examples. [Top] Collinear Gabors form a vertical contour (right of center), with randomly oriented flankers. [Bottom Left, Bottom Right] Same, but with the contours to the left of center, and with parallel or orthogonal flankers. Flanker position is bracketed in each panel with orientation distributions noted above. The bottom left and right panels are zoomed in for detail. Adapted from Figure 1 in Schallmo, Sponheim, & Olman (2013).

### Procedure

Participants were instructed to fixate on a cross at the center of the monitor and use their peripheral vision to detect laterality of the target contour within the stimuli grid. Responses were indicated via a left or right press on a button box. Degree of collinearity within target contours was jittered in increments of 4.5° with a floor of 0° and ceiling of 45°. Before the EEG session, participants completed a preliminary version of the Collinear Gabor Contour Task (CGCT) to determine collinear jitter threshold values for the EEG version of the task. In this preliminary task, collinear jitter was increased after three correct responses and decreased after one incorrect response. This staircase procedure was implemented to adjust task difficulty such that participant’s overall accuracy approached 79% (see (Schallmo *et al.* 2013) for more details). Setting jitter thresholds for each participant allowed for personalized task difficulty and prevented performance effects (i.e., floor effects and ceiling effects) during the EEG task. Additionally, individually tailored jitter thresholds helped control for effects related to generalized cognitive deficits in schizophrenia because task difficulty was equated across subjects (Gold & Dickinson 2013).

### EEG data collection and analyses

EEG collection and analysis information can be found in the supplemental materials. Stimulus locked event-related potentials (ERPs) were computed by averaging trials within each flanker condition for each participant. In order to differentiate neural responses associated with hypothesized temporally discrete mechanisms of surround suppression (Webb *et al.* 2005; Schallmo & Murray 2016), our ERP components of interest included earlier P1 (70-110ms) and N1 (110-210ms) components, and later P2 (190-290ms) and P3 (350-650ms) components. Time windows of interest were identified via inspection of grand average butterfly plot waveforms, grand average topographies, and histograms depicting mean amplitude frequency across subject-level ERPs. Electrodes where ERP amplitudes were largest or task effects were most evident, collapsed across groups, were selected for quantification. PO7 and PO8 electrode sites were averaged together for trials in which the target was contralateral to electrode location and quantified for P1, N1 and P2. Additionally, a late positive (P3) component was observed and quantified at CPz (350-650ms).

### Analysis

Subjects were excluded from analysis if accuracy scores were lower than 60% across all trials or lower than 75% on catch trials. Catch trials were trials in which the orientation jitter was 0° (i.e., the jitter level with the lowest difficulty). Contour detection thresholds were calculated by averaging contour jitter levels across trials within each flanker condition. Contextual modulation indices were then calculated by subtracting random condition jitter thresholds from both parallel and orthogonal jitter thresholds (Schallmo *et al.* 2013).

We ran eight separate repeated measures ANOVAs with group and gender as the between-subjects factors, flanker condition as the within-subjects factor and accuracy, response time, jitter threshold, contextual modulation index, P1 amplitude, N1 amplitude, P2 amplitude and P3 amplitude as the dependent variable, respectively. Contextual modulation indices were selectively correlated with corresponding P2 and P3 subtraction amplitudes to examine the significance of ERP modulation effects (i.e., random amplitudes were subtracted from orthogonal and parallel amplitudes to mirror the calculation of contextual modulation indices). All *p* values were corrected for multiple comparisons using Tukey’s HSD for between-subjects effects and False Discovery Rate (FDR; (Benjamini & Hochberg 1995) for within-subjects effects when appropriate).

## Results

### Task Performance

Contour detection performance was calculated by averaging jitter thresholds (degree of misalignment tolerated for the five Gabor elements that make up the linear contour) in each flanker orientation condition for each subject. There were no group differences in accuracy (main effect of group, ANOVA, *F*(3,102) = .91, *p* =.44) reflecting the individualization of difficulty level as part of the task. Average reaction times did not differ across groups (main effect of group, ANOVA, *F*(3,102) = 1.62, *p* =.19), but females exhibited slower reaction times compared to males (main effect of gender, ANOVA, *F*(1,102) = 4.70, *p* =.03).

Figure 2 depicts behavioral performance on the CGCT for each of the four groups. Higher jitter thresholds indicated stronger contour integration and better overall performance. We observed differences in contour detection performance between flanker conditions (relative orientation of Gabor elements neighboring the collinear contour: parallel, random, orthogonal) and groups (main effect of flanker, *F*(2,101) = 93.81, *p* < .001; main effect of group, *F*(3,102) = 4.00, *p* =.01). Irrespective of group, participants exhibited less tolerance to orientation jitter (i.e., worse contour detection performance) during the parallel flanker condition as compared to the random (FDR corrected *p* < .001) and orthogonal conditions (FDR corrected *p* <.001). Differences in contour detection between the random and orthogonal conditions were also significant (FDR corrected, *p* = .043).

**Figure 2.**
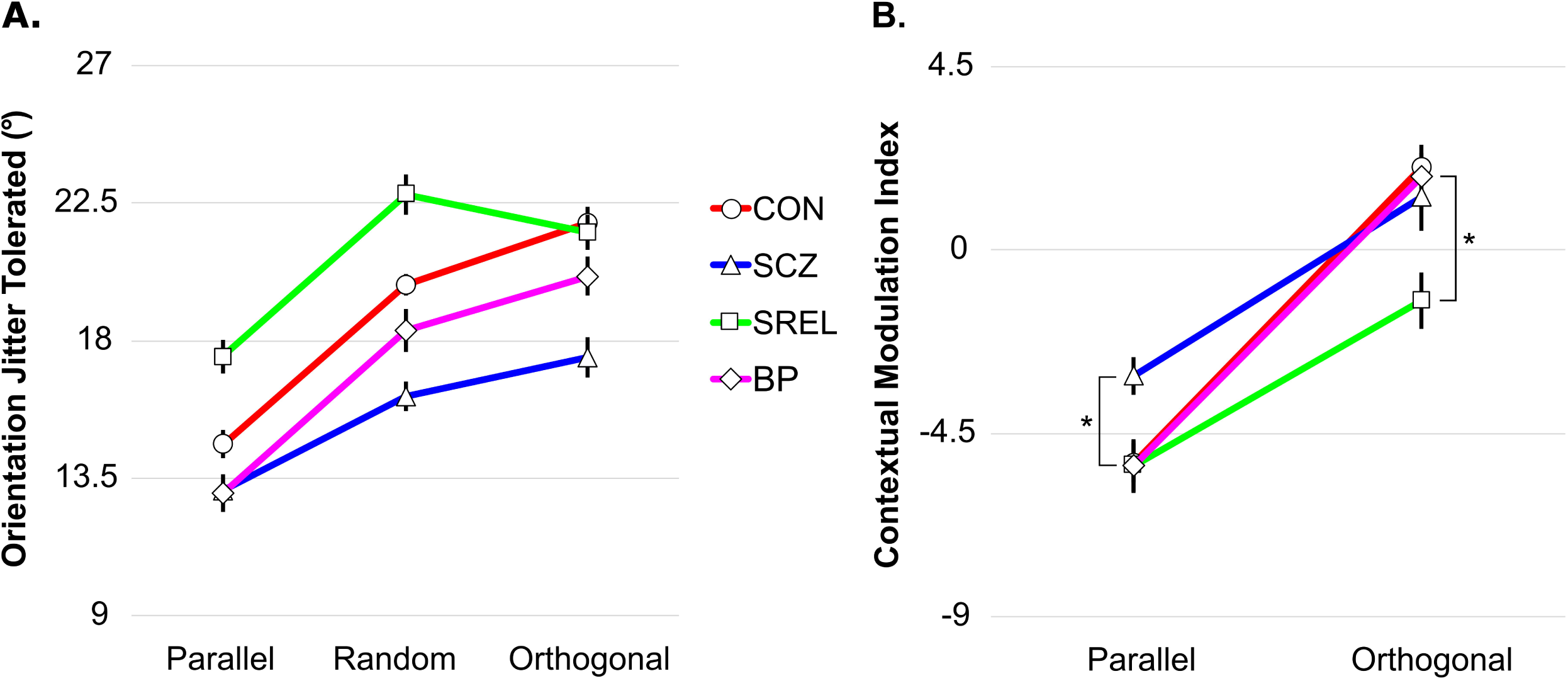
Contour Detection Thresholds and Contextual Modulation Indices. Mean detection thresholds [Left] and contextual modulation indices [Right] are plotted for 37 CON (circles), 27 SCZ (triangles), 23 SREL (squares) and 23 BP (diamonds) for Parallel, Random, and Orthogonal conditions. Asterisks denote attenuated suppression by parallel flankers in SCZ as compared to other groups (FDR corrected *p*s < .016) and attenuated facilitation in SREL as compared to CON (FDR corrected *p* = .018). Error bars are within-subjects standard error of the mean with a Morey correction factor. These were calculated by subtracting the within-subject mean across conditions from each subject’s data, and then adding the grand mean (across subjects and conditions), according to an established method (Morey 2008). This was done to help visualize variability across conditions, while accounting for the greater variability across individuals in ERP amplitudes.

Additionally, the effect of flanker orientation on contour detection differed between groups as indicated by an interaction of condition and group (*F*(6,204) = 3.35, *p* = .004). Across conditions, SCZ had lower jitter thresholds than SREL (Tukey HSD, *p* < .001) and CON (Tukey HSD, *p* = .016), but did not significantly differ from BP (Tukey HSD, *p* = .547). Notably, only CON (FDR corrected *p* =.004) exhibited significant facilitation of contour perception in the orthogonal condition as compared to the random condition.

### Contextual Modulation Performance

Contextual modulation indices were calculated to reflect the relative impact of flanker orientation on contour integration relative to the random flanker condition. As expected, we observed differences in contour detection between parallel-random and orthogonal-random modulation indices (main effect of relative flanker orientation, ANOVA, *F*(1,102) = 140.99, *p* < .001). Given our *a priori* hypothesis that SCZ would exhibit weakened contextual suppression (Schallmo *et al.* 2013; Mittal *et al.* 2015), we conducted one-tailed post hoc *t*-tests to assess SCZ performance in the parallel condition as compared to other groups. Analyses revealed that SCZ exhibited weaker modulation (i.e., less suppression) to parallel contextual stimuli as compared to the other groups (FDR corrected *p*s < .016). Unexpectedly, SREL exhibited negative estimated orthogonal modulation indices, signifying a lack of facilitation by orthogonal flankers. Follow-up pairwise comparisons revealed differences in contextual facilitation between SREL and CON only (FDR corrected *p* = .036). Interestingly, group differences in contextual modulation were affected by gender irrespective of flanker condition (interaction of group and gender, ANOVA, *F*(3,102) = 3.86, *p* = .012). This interaction was primarily driven by BP in which males exhibited more positive contextual modulation indices, collapsed across conditions, than females (FDR corrected *p* = .001).

### Visual Evoked Potentials: P1 and N1

We did not observe effects of group (ANOVA, *F*(3,109) = 1.6, *p* = .16), condition (ANOVA, *F*(2,109) = .4, *p* = .67) or gender (ANOVA, *F*(1,109) = .97, *p* = .33) on P1 mean amplitude at sites PO7/PO8. Furthermore, amplitude did not significantly differ across groups as a function of flanker orientation (interaction of group and condition, ANOVA, *F*(6,109) = .56, *p* = .76). Given the apparent group differences at P1 in the grand average waveforms (see Figure 3), we conducted follow up pairwise comparisons. This exploratory analysis revealed an attenuation of P1 mean amplitude in SCZ as compared to CON (uncorrected *p* = .034). Otherwise, P1 mean amplitude did not appear to modulate meaningfully as a result of task manipulations or psychopathology.

**Figure 3.**
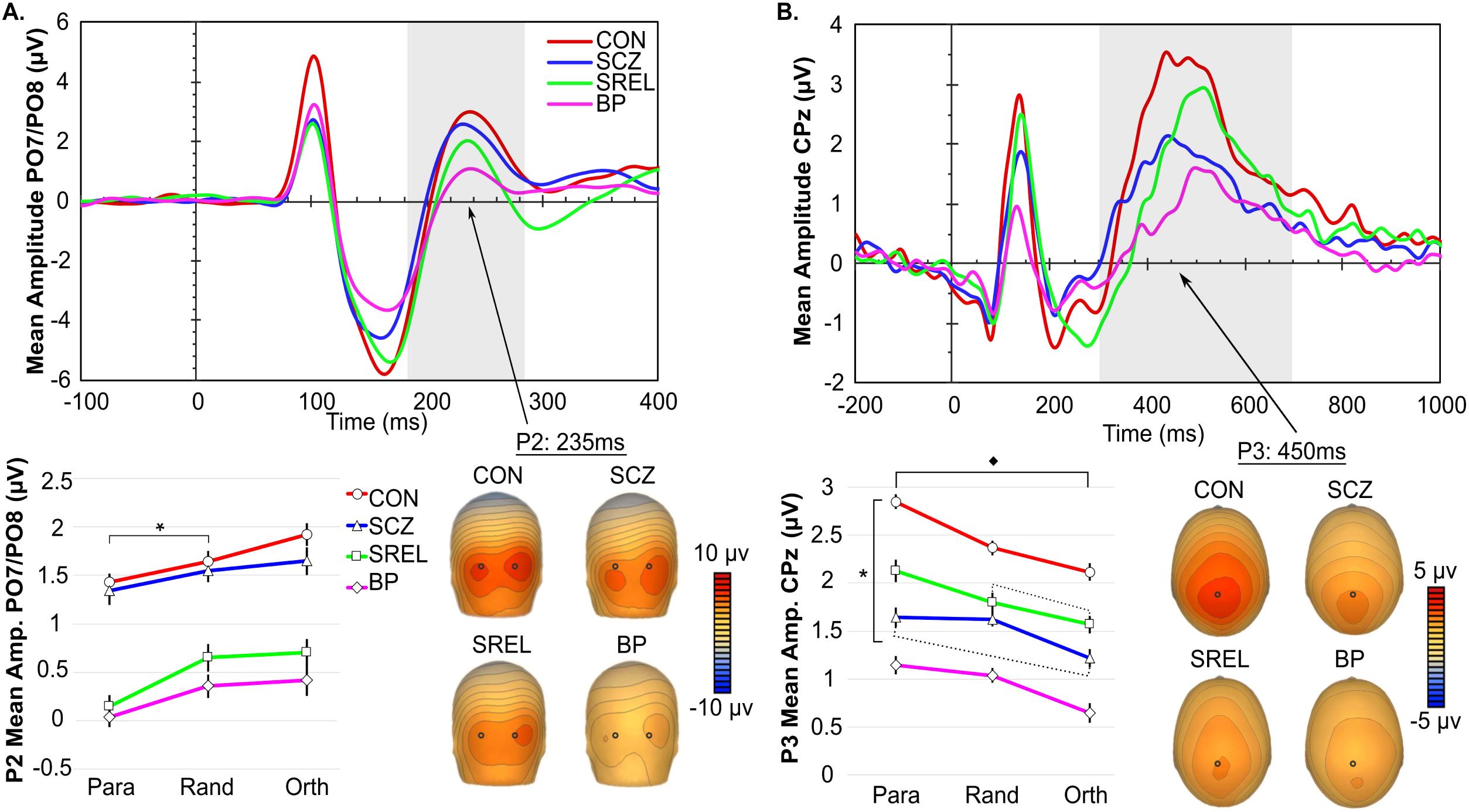
P2 Component and P3 Components. [Panel A and B, top] Grand average PO7/PO8 and CPz waveforms for each group. [Panel A and B, bottom left] P2 and P3 mean amplitudes with flanker condition on the x-axis. [Panel A and B, bottom right] Topographical representation of occipital P2 and centroparietal P3 activation for each group. [Panel A] * indicates significant amplitude modulation across all groups between parallel and random flanker conditions (FDR corrected *p*s <.002). [Panel A] Dotted lines indicate a lack of modulation between Parallel and Orthogonal conditions for SCZ and BP only. [Panel B] Black diamond indicates all groups significantly modulated P3 amplitude between parallel and orthogonal conditions except for SCZ (FDR corrected *p* =.137). [Panel B] * indicates lower overall P3 amplitudes in BP and SCZ compared to CON (Tukey’s HSD, *p*s < .019). [Panel B] Dotted lines indicate a lack of modulation between Parallel and Orthogonal flankers for SCZ, and a lack of modulation between Random and Orthogonal conditions for SREL. Error bars are within-subjects standard error of the mean with a Morey correction factor (see Figure 2). Note: we did not quantify the earlier positive component apparent in the CPz grand average waveform because inspection of grand average topographies revealed this component to be the result of a dipole generated by the occipital N1 component.

For N1 at sites PO7/PO8, we found a significant effect of flanker orientation on mean amplitude (main effect of flanker condition, ANOVA, *F*(2,101) = 3.31, *p* = .041). Post hoc pairwise comparisons revealed a difference in amplitude between parallel and orthogonal conditions (uncorrected *p* =.024). We did not observe differences in mean amplitude between groups (main effect of group, ANOVA, *F*(3,102) = .76, *p* = .52) or differences in the effect of flanker condition between groups (interaction of flanker condition and group, ANOVA, *F*(6,204) = .77, *p* = .59).

### Late Visual Event-Related Potential: P2

Figure 3 depicts visual ERP data from the CGCT that peaked over occipital brain regions. Main effects of flanker condition (ANOVA, *F*(2,101) = 9.80, *p* < .001) and group (ANOVA, *F*(3,102) = 3.22, *p* = .026) were found for the P2 component at sites PO7/PO8. Collapsed across groups, P2 amplitudes differed between parallel and random conditions (FDR corrected *p* = .002), and between parallel and orthogonal conditions (FDR corrected *p* = .002), but did not significantly modulate between random and orthogonal conditions (FDR corrected *p* = .504). This lack of modulation between random and orthogonal conditions is in line with behavioral findings in which the weakest flanker effects on contour detection were observed between random and orthogonal flanker conditions. Follow up pairwise comparisons of the group effect only revealed attenuated P2 amplitudes in BP compared to CON that did not withstand correction for multiple comparison (uncorrected *p* = .027). Although the effect of flanker condition on mean amplitude did not differ between groups (interaction of condition by group, ANOVA, *F*(6,204) = .50, *p* = .807) an exploratory, follow-up analysis of the simple main effects of condition on group showed that CON (FDR corrected *p* = .036) and SREL (FDR corrected *p* = .044) modulated P2 amplitudes between parallel and orthogonal conditions while SCZ and BP did not (FDR corrected *ps* >.05).

### Cognitive Event-Related Potential: P3

Figure 4 depicts ERP data from the CGCT that peaked over central brain regions. For the P3 component at scalp site CPz, we observed main effects of flanker (ANOVA, *F*(2,101) = 19.356, *p* < .001) and group (ANOVA, *F*(6,102) = 7.64, *p* < .001), but did not observe an interaction of group by flanker condition (ANOVA, *F*(6,204) = 1.03, *p* = .408). Collapsed across groups, participants showed the largest P3 amplitudes for contours with Parallel flankers, smallest P3 amplitudes for Orthogonal stimuli, whereas P3 responses in the Random condition were intermediate (FDR corrected *p*s < .002). Irrespective of condition, SCZ (Tukey’s HSD, *p* = .019) and BP (Tukey’s HSD, *p* = .001) exhibited reduced P3 amplitudes as compared to CON. Exploratory analysis of the simple main effects of condition on group revealed that SCZ failed to modulate P3 amplitude between the orthogonal and parallel condition (FDR corrected *p* =.137) while all other groups did (FDR corrected *ps* < .05). Additionally, SCZ and SREL failed to modulate P3 mean amplitude between the random and orthogonal conditions (FDR corrected *ps* >.05) in contrast with BP (FDR corrected *p* =.041) and CON (FDR corrected *p* =.038). This lack of modulation in SCZ and SREL aligns with the behavioral findings in which SCZ and SREL showed a lack of facilitation in contour detection between random and orthogonal conditions. No significant effects of gender were observed in any of the ERP analyses with the exception of a significant P3 group by gender interaction (ANOVA, *F*(3,109) = 2.89, *p* = .039). Follow-up pairwise comparisons revealed this interaction to be driven by SREL in which females exhibited larger P3 amplitudes than males (FDR corrected *p* = .011).

**Figure 4.**
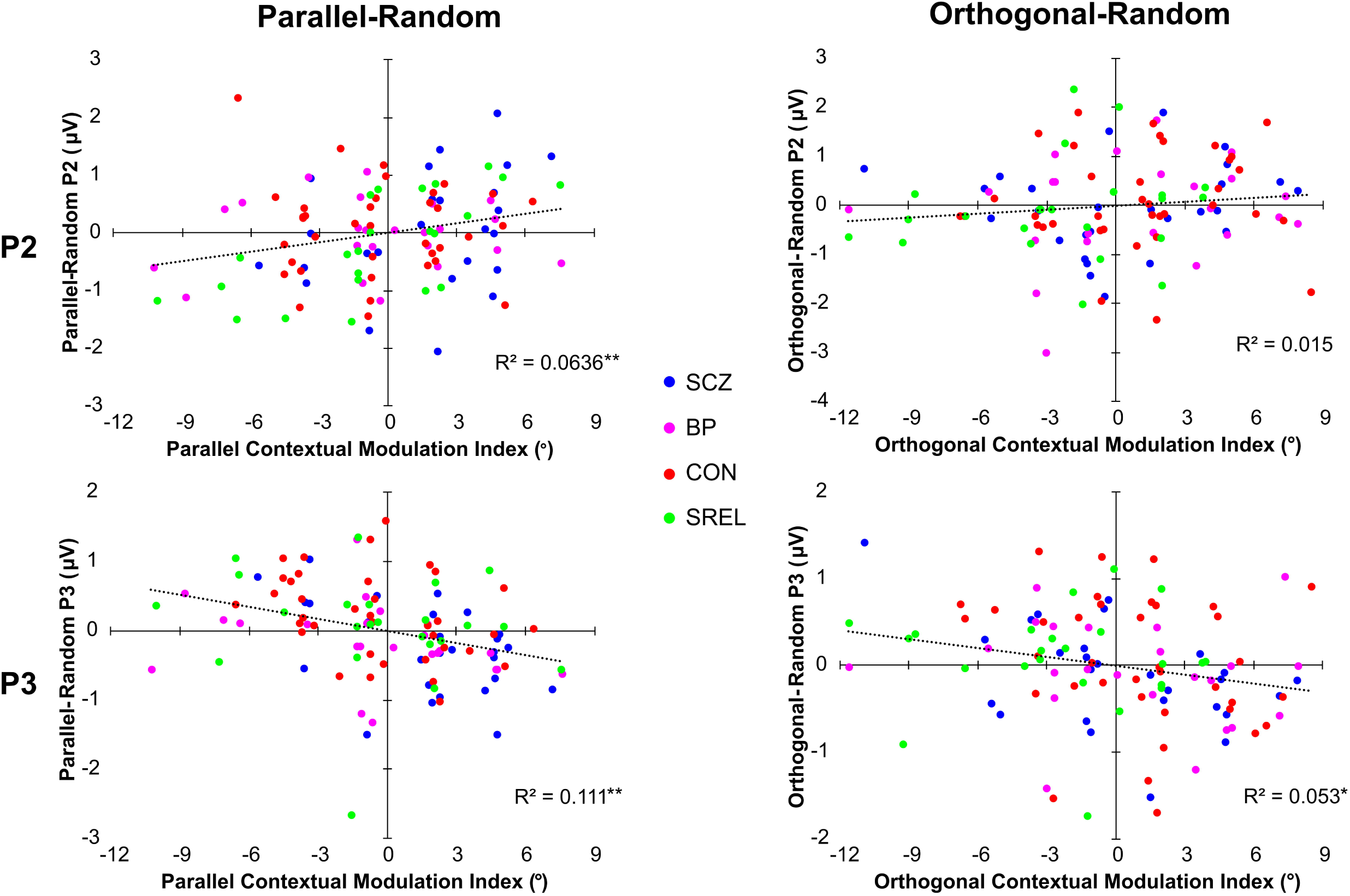
Associations Between Behavioral and ERP Contextual Modulation Indices. Scatterplots of P2 and P3 mean amplitude modulation indices and behavioral contextual modulation indices: CON (red), SCZ (blue), SREL (green), BP (pink). The trend line for each plot is calculated across groups. *R*^2^ values with a single asterisk represent significance when alpha is set at .05 while two asterisks represent significance when alpha is set at .01.

### Relationship of Contextual Effects on Contour Perception and ERP Responses

P2 and P3 amplitudes evoked in the random flanker condition were subtracted from orthogonal and parallel condition amplitudes in order to create ERP-based contextual modulation indices. These ERP contextual modulation indices were examined for associations with behavioral measures of flanker modulation of contour perception from the CGCT. Figure 5 depicts the relationships between perceptual and ERP modulations indices. Pearson correlations indicated a significant positive relationship between parallel-random P2 amplitudes and parallel-random modulation indices (*r*(109)=.252, FDR corrected *p* =.016), but did not indicate a significant relationship between orthogonal-random amplitudes and orthogonal-random modulation indices (r(109)=.122, FDR corrected *p* =.203). Similar correlations were calculated for P3 amplitudes which revealed a significant negative relationship between parallel-random P3 mean amplitudes and parallel contextual modulation indices (r(109)=-.333, FDR corrected *p* =.004), as well as orthogonal-random P3 mean amplitudes and orthogonal contextual modulation indices (r(109)=-.230, FDR corrected *p* =.021). Importantly, correlations with the perceptual modulation indices were in the opposite direction for the P2 and P3 ERP responses. Finally, we correlated ERP and behavioral contextual modulation indices with clinical symptom ratings (BPRS, SPQ, SGI), but these correlations did not reach significance (FDR corrected *p*s*>.05)*.

## Discussion

### Summary

In the present study, we administered a contour integration/contextual modulation task to patients with schizophrenia, their first degree biological relatives, patients with bipolar disorder, and healthy controls. We collected and analyzed simultaneously recorded behavioral and ERP data in an attempt to better understand the neurobiology of abnormal visual perception in psychotic disorders. Behavioral data were indicative of impaired contour integration (Silverstein & Keane 2011), and weakened contextual suppression in SCZ (Schallmo *et al.* 2013). Furthermore, BP exhibited intermediate contour integration impairments, but not contextual processing impairments. These findings strengthen theories that weakened surround suppression is specific to schizophrenia and not a product of broad cognitive impairment or general psychopathology (Tibber *et al.* 2013).

Additionally, we replicated previous findings in which SREL exhibited normal contour integration performance (Schallmo *et al.* 2013), but, in contrast to our previous findings, we observed that SREL exhibited abnormal contextual facilitation performance (i.e., for orthogonal vs. random flankers). These results suggest that SCZ and SREL may share some features of aberrant contextual modulation, but contour integration itself is spared among SREL. Interestingly, SREL exhibited normal contextual suppression (i.e., for parallel vs. random flankers) unlike SCZ, suggesting a subtle distinction in abnormal contextual modulation between SCZ and SREL.

EEG acquired during task performance provided a direct measurement of neural activity related to surround suppression and contour integration. The paucity of significant effects of group or condition for P1 and N1 components suggests that differences in task performance were not strongly reflected in brain activity within 0-200 ms after stimulus presentation. Instead, robust flanker and group effects in the P2 and P3 components suggest that deficits in surround suppression and contour integration are in part related to higher order perceptual processing (Keane *et al.* 2014; Silverstein *et al.* 2015b). Moreover, we observed that P2 and P3 amplitudes were correlated with behavioral contextual modulation indices such that individuals for whom contour perception was more strongly affected by flanker orientation also showed greater P2 and P3 amplitude differences between flanker conditions. These associations were positive for P2 and negative for P3 signals, reflecting the opposite pattern of response amplitudes across flanker conditions between the two brain responses. These correlations suggest an association between neural responses between 200-600 ms post-stimulus and the effect of surrounding context on the perceptual salience of visual contours. Notably, these results expand upon the ERP findings of Butler et al., 2013; in addition to reporting a similar set of ERP abnormalities, we show that such abnormalities are not dependent on higher-order object identification processes (as they are present for simple linear contours), implying the extensive effects of aberrations in simple perceptual processes.

Reduced P3 amplitudes at centroparietal midline sites in SCZ and BP, as compared to CON, are consistent with previous findings (Jeon & Polich 2003; van der Stelt *et al.* 2004; Bramon *et al.* 2005; Luck *et al.* 2009; Ryu *et al.* 2010; Johannesen *et al.* 2013; Maekawa *et al.* 2013), however further research is needed to clarify observed changes in P3 amplitude between contextual conditions. Literature suggests that P3 amplitude is affected by a variety of factors including frequency of stimulus presentation (e.g., rare vs. common), stimulus/task complexity, information transmission and context updating (Johnson 1986; Donchin & Coles 1988). Most relevant to our task is the premise of information transmission. According to Johnson (1986), “changes in P300 amplitude that are accompanied by changes in the subject’s level of performance are, by definition, due to changes in the amount of transmitted information.” Thus, differences in P3 amplitude may reflect differences in perceived contour salience (i.e., differing levels of information transmission) across the three flanker conditions in our task (Kok 2001).

Alternatively, there is a growing body of literature investigating the P3 (also referred to as the centroparietal positivity [CPP] in perceptual literature) as a supramodal, higher order evidence accumulation mechanism (O’Connell *et al.* 2012; Kelly & O’Connell 2013; Twomey *et al.* 2015; Tagliabue *et al.* 2019). Importantly, Tagliabue et al. (2019), reported that the CPP is dependent upon subjective perceptual experience (top-down processing) even after controlling for objective stimulus intensity. In the CGCT, SCZ were the only group that did not significantly modulate P3 amplitude between flanker conditions which, given the CPP literature, may suggest aberrant high-level processing in SCZ. Viewing the P3 in the context of subjective sensory accumulation may help explain observed opposing CGCT behavioral associations with P2 and P3 amplitude. Specifically, it is possible that the P2 reflects higher order visual processing in the occipital cortex while P3 reflects information accumulation in the parietal cortex. Thus, in the CGCT, smaller and larger P2 responses may reflect contextual suppression and facilitation respectively, while smaller and larger P3 amplitudes may reflect less vs. greater sensory evidence accumulation respectively.”

To summarize, the current study provided novel neurophysiological correlates of well-documented visuoperceptual abnormalities in SCZ. These findings suggest that abnormal contour integration and contextual modulation in SCZ likely relate to and may result in aberrant high-level visual cortical and cognitive functions. By examining BP and SREL, we probed the extent to which deficits in contour integration and contextual suppression are specific to SCZ. We found a more subtle deficit in contour detection performance in BP subjects as compared to SCZ, that was reflected in intermediate and late latency ERPs (P2 and P3). SREL subjects showed generally intact patterns of neural responses during visual contour perception, but did exhibit abnormally weak orthogonal facilitation of contour detection in our behavioral task. Together, our findings point to specific neural deficits in later-stage visual processing in SCZ.

## Acknowledgements

The authors would like to thank current and past members of the Cognition and Brain Sciences lab at the University of Minnesota for fostering a welcoming community devoted to excellence in research. Specifically, we would like to thank Collin Teich, Josh Stim and Dr. Seung Suk Kang for their help with preprocessing these EEG data and scripting in Matlab.

## Ethical Standards

The authors assert that all procedures contributing to this work comply with the ethical standards of the relevant national and institutional committees on human experimentation and with the Helsinki Declaration of 1975, as revised in 2008.

