## Supplemental Text for "Reduced Influence of Perceptual Context in Schizophrenia: Behavioral and Neurophysiological Evidence"

Supplemental Information

Methods

*Exclusion Criteria*

SCZ, BP and CON were excluded from the study based upon the following criteria: English as a second language, age > 60 years, IQ < 70, poor visual acuity that could not be corrected to a logarithmic visual acuity score of < 0.3 (LIGHTHOUSE Distance Visual Acuity Test, Long Island City, NY), substance dependence in the past six months, substance abuse within two weeks of testing, history of head injury with skull fracture or substantial loss of consciousness, electroconvulsive therapy, amblyopia uncorrected before age 18, epilepsy, stroke, or other neurological conditions. Additional exclusion criteria for CON were personal or immediate family history of bipolar disorder, major depressive disorder, or a psychotic disorder. SREL were excluded only if they had poor visual acuity that could not be corrected to normal (i.e., logarithmic visual acuity score of < 0.3), or a medical condition that prevented participation.

*Stimuli*

Stimuli for the CGCT were presented using E-Prime on a Dell computer running Windows XP. The images were displayed on a NEC 17" CRT monitor with a resolution of 1024 x 768. Monitor viewing distance was 61cm and the visual angle subtended 35.1 x 26.7 degrees. Stimuli consisted of Gabor patches organized into 15 x 15 grids (see Figure 1). The orientation of orthogonal flanker stimuli ranged from 45° to 135° such that the mean orientation of flankers was perpendicular to the target contour. Similarly, orientation of parallel flanker stimuli ranged from -45° to 45° such that the mean orientation of flankers was parallel to the contour. Random flanker stimuli ranged from -90° to 90°. Non-contour, non-flanker gabor patches were randomly oriented, but cardinal neighbors differed by at least 30° to prevent stochastic contour formation.

*Procedure*

In both versions of the task (pre-EEG and EEG), participants completed one practice block consisting of 20 trials followed by four, 60 trial blocks with pseudorandomized flanker conditions. Breaks between blocks were not timed and all blocks were initiated by the task administrator after verbally checking-in with the participant. The EEG version of the task was identical to the preliminary version except jitter levels were held constant during the EEG unless accuracy in any condition fell below 60%. In such cases, jitter was reduced by one step (i.e., 4.5°) for that condition.

At the beginning of each trial, a fixation cross appeared for 1000ms. Gabor stimuli were then presented for 250ms followed by a jittered response window of 500, 600, 700, 800, 900, or 1000ms. Following the response window, a 250ms feedback cue was shown in which a green fixation cross signified a correct response, a red fixation cross signified an incorrect response and a white fixation cross signified a missed response. The interstimulus interval ranged from 2000-2500ms depending on the duration of the response window. For each trial, the fixation point was randomly moved within .5° of the center to prevent participants fixating on potential target contour locations. Accuracy and reaction times were collected for all trials.

*EEG data collection and analyses*

EEG data were collected using the BioSemi ActiveTwo system with a differential amplifier and a high density 128 electrode cap with a radial layout system. The amplifier utilized a fixed first order analog anti-aliasing filter with a 3.6 kHz half power cut-off frequency (-3dB) and a 6dB/octave roll off. All channels were referenced to linked-ears during acquisition. Data were recorded with a sampling rate of 1024 Hz, and downsampled to 256 Hz offline with a high pass filter of .5 Hz and low pass filter of 256 Hz (both filters were 3rd order butterworth with an 18dB/octave roll off; frequency cutoffs were half power) . Malfunctioning electrodes (i.e., electrodes that were significantly uncorrelated with neighboring electrodes) were identified by visual inspection and replaced via spherical spline interpolation. The data were then epoched from -500 to 1500 ms relative to onset of the Gabor stimuli. Non-neural electrical activity including ocular, cardiac, muscular, and electrical noise were identified by applying independent component analysis (ICA) to the epoched data. Independent components (ICs) were then visually inspected and labeled; only ICs reflecting neural activity were included in the final reconstituted data. Denoised data were then re-referenced to the average scalp signal from all electrodes. Participants were excluded if they had less than 20 viable trials per condition after preprocessing. On average, each group had a similar number of trials per condition (SCZ, $M$ = 52.2; BP, $M$ = 51.5; SREL, $M$ = 51.2; CON, $M$ = 51.2; ANOVA, $F(3,106) = .56$ $p = .64$). ERPs were created with a 200ms prestimulus baseline correction, and a 30 hz lowpass filter (3rd order butterworth with an 18dB/octave roll off; frequency cutoff was half power).
